# Temporal enzymatic treatment to enhance the remodelling of multiple cartilage microtissues into a structurally organised tissue

**DOI:** 10.1101/2022.09.07.506986

**Authors:** Ross Burdis, Xavier Barceló Gallostra, Daniel J. Kelly

## Abstract

Scaffold-free tissue engineering strategies aim to recapitulate key aspects of normal developmental processes as a means of generating highly biomimetic grafts. Cartilage and fibrocartilaginous tissues have successfully been engineered by bringing together large numbers of cells, cellular aggregates or microtissues and allowing them to self-assemble or self-organize into a functional graft. Despite the promise of such approaches, considerable challenges still remain, such as engineering scaled-up tissues with predefined geometries, ensuring robust fusion between adjacent cellular aggregates or microtissues, and directing the (re)modelling of such biological building blocks into a unified scaled-up graft with hierarchical matrix organisation mimetic of the native tissue. In this study, we first demonstrate the benefits of engineering cartilage *via* the fusion of multiple cartilage microtissues compared to conventional scaffold-free approaches where (millions of) individual cells are allowed to aggregate and generate a cartilaginous graft. Key advantages include the engineering of a tissue with a richer extracellular matrix, a more hyaline-like cartilage phenotype and a final graft which better matched the intended geometry. A major drawback associated with this approach is that individual microtissues did not completely (re)model and remnants of their initial architectures where still evident throughout the macrotissue. In an attempt to address this limitation, the enzyme chondroitinase ABC (cABC) was employed to accelerate structural (re)modelling of the engineered tissue. Temporal enzymatic treatment supported robust fusion between adjacent microtissues, enhanced microtissue (re)modelling and supported the development of a more biomimetic tissue with a zonally organised collagen architecture. Additionally, we observed that cABC treatment modulated matrix composition (rebalancing the collagen:glycosaminoglycans ratio), tissue phenotype, and to a lesser extent, tissue mechanics. Ultimately, this work demonstrates that microtissue self-organisation is an effective method for engineering scaled-up cartilage grafts with a pre-defined geometry and near-native levels of ECM accumulation. Importantly we have demonstrated that key limitations associated with tissue engineering using multiple cellular aggregates, microtissues or organoids can be alleviated by temporal enzymatic treatment during graft development.

## Introduction

Engineering articular cartilage that mimics both the structure and composition of the native tissue remains a considerable challenge. Shortcomings associated with traditional top-down tissue engineering approaches has motivated interest in scaffold-free strategies that aim to mimic developmental processes to generate a more biomimetic tissue [1–9]. These approaches often yield highly biomimetic engineered cartilage as the cells, unconstrained by an interstitial scaffold material, are allowed to interact with and remodel the extracellular matrix (ECM) in a fashion that recapitulates key events in the native tissue’s developmental programme [10–12]. In an attempt to scale-up the engineering of more complex tissues and organs, emerging scaffold-free strategies seek to use cellular spheroids, microtissues, and/or organoids as biological building blocks that can be combined to fabricate larger regenerative implants. Such approaches have been applied to multiple tissues, including bone [13–17], cartilage [18,19], vasculature [15,20–24], osteochondral [25–29], and liver [30]. Compared to more traditional scaffold-free approaches, that use single cells as the minimal unit building block for generating an engineered tissue, biofabrication using cellular aggregates or microtissues as biological building blocks can offer several benefits [6]. For example, increasing the number and/or size of the biological building-blocks can facilitate the generation of larger tissues approaching those required for clinical use. Moreover, creating tissues/grafts with a predesigned geometry using individual cells can be challenging, requiring complex biofabrication strategies which could limit a widespread uptake. In contrast, aggregate based approaches have been successfully employed to create scaled-up, functional tissues/grafts with user-defined geometries using both manual and automated biofabrication methods [13,18,31,32]. Specifically, in the field of cartilage tissue engineering we have demonstrated the capacity to manually bioassemble microtissues within a 3D printed polymer framework to generate a biphasic osteochondral implant for joint resurfacing [9], while others have leveraged novel biofabrication strategies (including bioprinting) to spatially organise cell spheroids, organoids, microtissues, and tissue strands for the generation of tissues/grafts with defined geometries and architectures [13,15,29,31,33–36].

Despite considerable progress in this field, ensuring robust fusion between multiple biological building blocks (e.g. cellular spheroids/microtissues) and subsequently directing their remodelling into a unified tissue with biomimetic organisation remains a key challenge [6,24]. In particular, pervasive tissue architectures tend to form within each microtissue and are apparent throughout the engineered macrotissue [13,15,26,29,37]. In order to engineer functional tissues, these initial architectures must be effectively remodelled to generate a tissue with native structural organisation. In the context of cartilage tissue engineering, recapitulation of the physiological stratification found in native articular cartilage (AC) is rarely reported. Although the use of instructive scaffolds that provide boundary conditions to the developing tissue have gone some of the way in addressing the challenge of generating a stratified AC [9,38–40], complete recapitulation of the zonal organisation within engineered cartilage tissues remains elusive.

In this study, we first sought to demonstrate the benefits of engineering scaled-up cartilage grafts *via* the fusion of multiple cartilage microtissues compared to traditional scaffold-free strategies where the construct is generated *via* the self-organisation of individual cells. We then sought to address two of the key challenges associated with the biofabrication of zonally stratified articular cartilage using multiple cartilage spheroids or microtissues, namely 1) ensuring robust fusion between adjacent microtissues, and 2) directing the remodelling of the initial microtissue architectures into zonally stratified tissue mimetic of articular cartilage. To this end, we explored the use of chondroitinase-ABC (cABC) treatment to help direct the fusion and remodelling of multiple cartilage microtissues into zonally stratified tissue. Similar catabolic enzymatic regimes have been successfully employed to enhance the development of self-assembled cartilage in other scaffold-free systems [8,41–43]. Here we sought to determine the impact of such enzymatic treatment on tissue development during the self-organisation of multiple microtissues, with the aim of encouraging tissue (re)modelling and the generation of a cartilage graft with biomimetic composition and structural organisation.

## Materials and Methods

### Bone Marrow Mesenchymal Stem/Stromal Cell (MSC) Isolation and Expansion

MSCs were isolated from the femoral shaft of 4 month old pigs under sterile conditions. They were expanded in expansion medium (XPAN), which is composed of high glucose Dulbecco’s modified eagle’s medium (hgDMEM) GlutaMAX supplemented with 10 % v/v FBS, 100 U/mL penicillin, 100 μg/mL streptomycin (all Gibco, Biosciences, Dublin, Ireland) and 5 ng/mL FGF2 (Prospect Bio). All MSCs were expanded in physioxic conditions (37 °C in a humidified atmosphere with 5 % CO_2_ and 5 % O_2_) for chondrogenic differentiation. After initial colonies had formed, MSCs were trypsinised, counted, and re-seeded at a density of 5,000 cells/cm^2^ and expanded until the end of passage 2.

### Construct Self-organisation & Enzyme Treatment

#### Microwell Fabrication

The procedure for fabricating the hydrogel microwell platform and the method of generating cellular aggregates and microtissues has been described previously [14]. The same underpinning methodology was employed herein. Briefly, a novel stamp designed using Solidworks^®^ CAD software and fabricated using a Form 3 stereolithography (SLA) printer (Formlabs, Massachusetts, United States) was used as the positive mould for the microwell array. Positive moulds were processed post-printing in accordance with the manufacturer’s guidelines, cleaned and gas sterilised using ethylene oxide (EtO) prior to use (Anprolene^®^ gas sterilisation cabinet, Andersen Sterilizers). Non-adherent hydrogel microwells were fabricated under sterile conditions, within a class II biosafety cabinet. Firstly, agarose (Sigma Aldrich) was dissolved in phosphate buffered saline (PBS) (Sigma Aldrich) at a concentration of 4 % (w/v) and autoclaved to ensure sterility. The aforementioned sterile 3D printed positive moulds were then inserted into the agarose, ensuring no bubbles became trapped underneath the mould. Once cooled, excess solidified agarose above the mould was removed and the positive mould was pulled from the agarose, leaving the patterned agarose imprinted with the 401 microwells within each well. All agarose microwells were soaked overnight in XPAN prior to cell seeding.

#### Microtissue Generation and Macrotissue Engineering

Initially cartilage microtissues were formed using 3 × 10^3^ cells per microtissue. Chondrogenesis was initiated by culturing the cells in chondrogenic Differentiation Medium (CDM) for 2 days. CDM was made by supplementing hgDMEM GlutaMAX with 100 U/mL penicillin, 100 μg/mL streptomycin (both Gibco), 100 μg/mL sodium pyruvate, 40 μg/mL L-proline, 50 μg/mL L-ascorbic acid-2-phosphate, 4.7 μg/mL linoleic acid, 1.5 mg/mL bovine serum albumin, 1 × insulin–transferrin–selenium (ITS), 100 nM dexamethasone (all from Sigma), 2.5 μg/mL amphotericin B and 10 ng/mL of human transforming growth factor-β_3_ (TGF-β) (Peprotech, UK). After 2 days, microtissues were liberated from the microwells and harvested for future biofabrication steps.

The capacity form cartilage macrotissues *via* the spontaneous self-organisation of cartilage microtissues and single-cells was determined by seeding microtissues and a high-concentration single cell suspension into a custom agarose well. The well was created using sterile 2 % w/v agarose cast into a 12 well plate. The central agarose well was 3 mm in diameter and 1.5 mm in depth. The total number of cells seeded into the agarose well in both groups was 3 × 10^6^, meaning 3 × 10^6^ MSCs in a single cell suspension of 8 μL and 1 × 10^3^ microtissues each containing 3 × 10^3^ MSCs in 8 μL. After seeding the wells, the plate was centrifuged at 400 × g for 5 minutes to ensure the single-cells/microtissues were collected at the bottom of the well. Each macrowell was then topped up with 2 mL of chondrogenic medium and returned to the incubator and cultured in physioxic conditions (37 °C in a humidified atmosphere with 5 % CO_2_ and 5 % O_2_). After 7 days, microtissues were sufficiently fused to allow the removal of the macrotissues from the seeding well. The cartilage tissues were then cultured for the remainder of the study in a 12 well-plate coated with 2 % agarose to prevent cellular attachment.

#### Chondroitinase-ABC Treatment

On day 14, prior to enzymatic treatment, constructs were washed in hgDMEM. Following which, they were maintained in an enzymatic solution containing 2 U/mL cABC (Sigma-Aldrich), 0.05 M acetate (Trizma Base, Sigma-Aldrich) activator in hgDMEM for 4 hours at physioxic conditions. After the treatment, the engineered tissues were washed again with hgDMEM to ensure removal of any residual cABC, before the addition of fresh chondrogenic medium and the continuation of chondrogenic cultivation in physioxic conditions for the remaining 14 days.

### Biochemical Evaluation

Samples were washed in PBS after retrieval and the wet-weight was recorded immediately. An enzyme solution, 3.88 U/mL of papain enzyme in 100 mM sodium phosphate buffer/5 mM Na2EDTA/10 mM L-cysteine, pH 6.5 (all from Sigma–Aldrich), was used to digest the samples at 60 °C for 18 hours. DNA content was quantified immediately after digestion using Quant-iT™ PicoGreen ^®^ dsDNA Reagent and Kit (Molecular Probes, Biosciences). Sulphated glycosaminoglycan content (sGAG) was quantified with 1,9-dimethylene blue (DMMB) at pH 1.5; metachromatic changes of DMMB in presence of sGAG were determined using the Synergy HT multi-detection micro-plate reader (BioTek Instruments, Inc) at 530 and 590 nm. 530/590 absorbance ratios were used to generate a standard curve and determine sGAG concentration of unknown samples, chondroitin sulphate was used as standard (Sigma-Aldrich). Total collagen content was determined using a chloramine-T assay to measure the hydroxyproline content and calculated collagen content using a hydroxyproline-to-collagen ratio of 1:7.69 [44]. Briefly, samples were mixed with 38 % HCL (Sigma) and incubated at 110 °C for 18 hours to allow hydrolysis to occur. Samples were subsequently dried in a fume hood and the sediment reconstituted in ultra-pure H_2_O. 2.82 % (w/v) Chloramine T and 0.05 % (w/v) 4-(Dimethylamino) benzaldehyde (both Sigma) were added and the hydroxyproline content quantified with a trans-4-Hydroxy-L-proline (Fluka analytical) standard using a Synergy HT multi-detection micro-plate reader at a wavelength of 570 nm (BioTek Instruments, Inc).

### Histological & Immunohistochemical Evaluation

#### Histological Evaluation

Samples were fixed using 4 % paraformaldehyde (PFA) solution overnight at 4 °C. After fixation, samples were dehydrated in a graded series of ethanol solutions (70 % - 100 %), cleared in xylene, and embedded in paraffin wax (all Sigma-Aldrich). Prior to staining tissue sections (5 μm) were rehydrated. Sections were stained with haematoxylin and eosin (H&E), 1 % (w/v) alcian blue 8GX in 0.1 M hydrochloric acid (HCL) (AB) to visualise sulphated glycosaminoglycan (sGAG) content and counter-stained with 0.1 % (w/v) nuclear fast red to determine cellular distribution, 0.1 % (w/v) picrosirius red (PSR) to visualise collagen deposition, and 1 % (w/v) alizarin red (AR) (pH 4.1) to determine mineral deposition *via* calcium staining (all from Sigma-Aldrich). Stained sections were imaged using an Aperio ScanScope slide scanner and thickness measurements obtained using Aperio Imagescope.

#### Immunohistochemical Evaluation

Samples were fixed in 4% paraformaldehyde, dehydrated in a graded series of ethanol, cleared in xylene, and embedded in paraffin wax (all Sigma-Aldrich). Prior to staining, tissue sections (5 μm) were rehydrated. Antigen retrieval was carried out with pronase (3.5 U/ml; Merck) at 37 °C for 25 minutes, followed by hyaluronidase (4000 units/ml; Sigma-Aldrich) at 37 °C for 25 minutes for collagen type I and type II. For collagen type X, an initial treatment with pronase (35 U/mL; Merck) at 37 °C for 5 minutes, was followed by chondroitinase ABC (0.25 U/mL; Sigma-Aldrich) at 37 °C for 45 minutes. Non-specific sites were blocked using a 10% goat serum and 1% BSA blocking buffer for 1 hour at room temperature. Collagen type I (1:400; ab138492; Abcam), type II (1:400; sc52658; Santa Cruz), and type X (1:300; ab49945; Abcam) primary antibodies were incubated overnight at 4 °C, followed by 20 minutes treatment using 3% hydrogen peroxide solution (Sigma-Aldrich). Secondary antibodies for collagen type I (1:250; ab6720; Abcam), type II (1:300; B7151; Sigma-Aldrich) and type X (1:500, ab97228; Abcam) were incubated for 4 hours at room temperature. Samples were then incubated for 45 minutes with VECTASTAIN Elite ABC before staining them with ImmPACT DAB EqV (both from Vector Labs).

### Polarised-light Microscopy & Collagen Alignment Quantification

Sections stained with PSR were imaged using polarised light microscopy to visualise collagen fibre orientation. Quantification of mean fibre orientation and dispersion was carried out using the ‘directionality’ feature in ImageJ software. Fibre coherency and colour maps were determined using the OrientationJ plugin [45]. The zones of the engineered tissue were defined as follows: the deep-zone was characterised as the lower 50 % of the tissue, the middle-zone was the intermediate 40 %, and the superficial zone was the top ~10 % of the engineered tissue. Multiple sections were taken throughout the engineered samples. From these sections, 5 were selected randomly for quantification. Data points presented graphically represent the quantification of the fibre directionality from the defined zones of the engineered and native tissue. Specifically, orientation graphs show mean average and standard deviation from the histograms generated using the directionality feature in ImageJ software. 95 % confidence ellipses presented in dispersion vs orientation plots were determined using the Real statistics resource pack add-in for excel.

### Mechanical Evaluation

To investigate how tissue maturation and enzymatic treatment influence the mechanical properties of cartilages engineered *via* the self-organisation of early-cartilage microtissues, unconfined compressions tests were carried out in a PBS bath using a single column Zwick testing machine (Zwick, Roell, Herefordshire, UK) equipped with a 10 N load cell. To ensure contact between the surface of the constructs and the top compression platen, a preload of 0.05 N was used. A peak of 10 % strain was applied at a rate of 1 mm/min and the equilibrium stress was obtained after a relaxation time of 15 minutes. After the relaxation phase, five compression cycles at 1 % strain at a frequency of 1 Hz were superimposed. The ramp modulus was calculated as the slope of the initial linear region of the obtained stress-strain curves. The equilibrium modulus was determined as the equilibrium force divided by the sample’s cross-sectional area divided by the applied strain. The dynamic modulus was measured as the average force amplitude over the five cycles divided by the sample’s cross-sectional area divided by the applied strain amplitude.

### Statistical Analysis

Statistical analysis was performed using GraphPad Prism software (GraphPad Software, CA, USA). The statistical analysis performed in each case is described in the table/figure caption. Numerical and graphical results are presented as mean ± standard deviation (SD) throughout. Significance was determined as follows; ns (not significant), p > 0.05, * p < 0.05, ** p < 0.01, *** p <0.001, and **** p < 0.0001. Different symbols donating significance are used in places, but the level of significance is consistent throughout.

## Results

### Cartilage microtissues self-organise into a more hyaline cartilage-like tissue compared to single cells

Given there are limited examples of directly comparing single cell- and microtissue-based self-organisation strategies, this study first sought to elucidate which method yielded a superior *in vitro* cartilage. To investigate this, comparable numbers of cells (in either single-cell or a microtissue format) were placed into hydrogel microwells where neo-tissue growth and maturation was quantified over 28 days of chondrogenic cultivation, with weekly biochemical and histological evaluation (**Fig. 1**). In both groups, total levels of sGAG and collagen increased predictably throughout the duration of the culture (**Fig. 2A**). After 28 days, the cartilage engineered *via* the self-organisation of cartilage microtissues contained significantly higher levels of both sGAG and collagen compared to its single-cell counterpart (2 fold higher total sGAG and 1.2 fold higher total collagen), while the DNA content was comparable in both groups. The sGAG-to-collagen ratio within both engineered cartilages revealed a non-physiological bias towards a GAG rich ECM (**Fig. 2A**). Normalisation of the absolute amounts of sGAG to DNA quantity demonstrated that biosynthesis of sGAG by resident cells was significantly higher in the microtissue group at days 21 and 28. There was no significant differences in biosynthetic output of collagen per cell (**Fig. 2B**). By day 28, both groups exhibited near-native levels of sGAG as a percentage of tissue wet weight (4.879 ± 0.174 % and 5.546 ± 0.245 % for single cell and microtissues respectively). Despite this evidence of robust chondrogenesis, the levels of collagen (both absolute and normalised) are noticeably lower than those in native articular cartilage (**Fig. 2C**).

**Figure 1.**
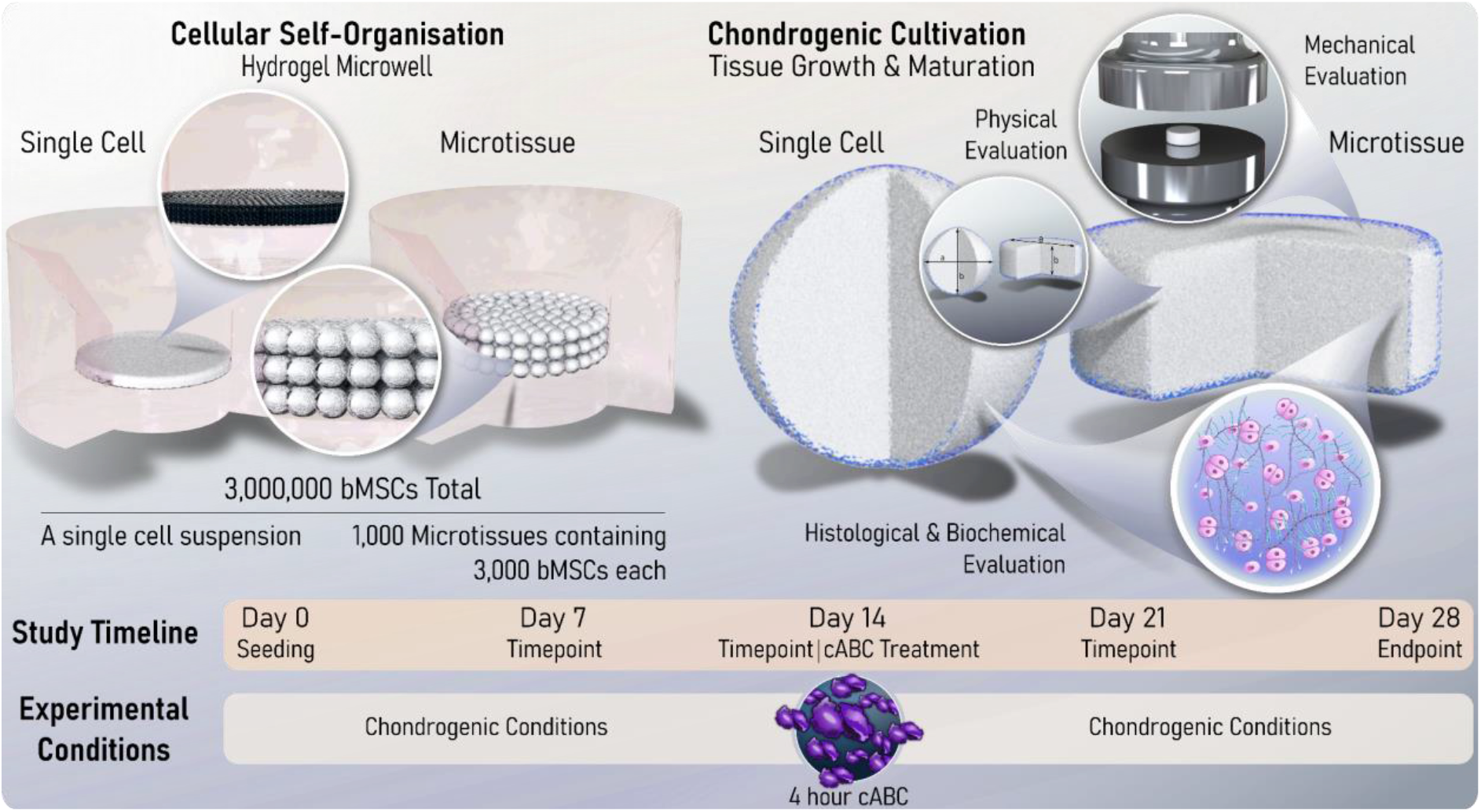
Study Schematic. Two different cellular self-organisation strategies are employed in this study. The first involves the self-organisation of 3 × 10^6^ bMSCs within a non-adherent hydrogel well. The second uses 1 × 10^3^ early-cartilage microtissues, each containing 3 × 10^3^ MSCs, as biological building blocks. A study timeline is also provided that indicates the weekly endpoints used to map tissue maturation. In one arm of the study, a 4 hour chondroitinase-ABC (cABC) treatment was undertaken on day 14. Finally, a schematic representation of the analytical techniques employed to determine the quality of the self-organised cartilages generated within this study is provided. Collectively, this work aims to provide a comprehensive timeline for tissue growth and maturation of self-organised cartilages, as well as determining the effect enzymatic treatment has on the quality of cartilage macrotissues bioassembled using single cells and cartilage microtissues as biological building blocks.

**Figure 2.**
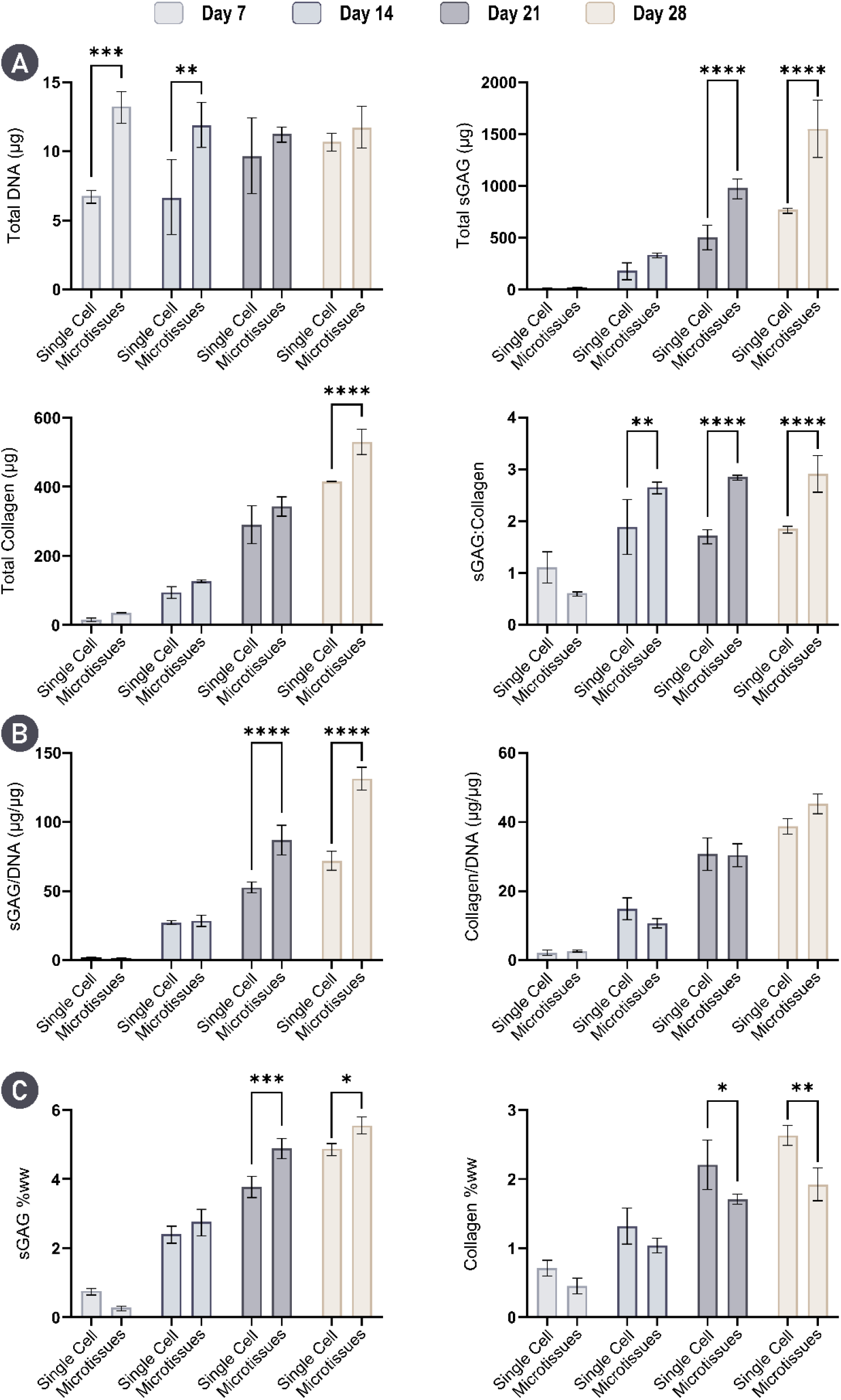
Cartilage microtissues self-organise a cartilage-rich ECM. A) Total levels of key cartilage ECM components; DNA, sGAG, and collagen are provided at weekly timepoints for 28 days of chondrogenic cultivation. Total levels of sGAG and collagen are significantly higher in the microtissue group by day 28. Additionally, the sGAG to collagen ratio within the engineered tissues is given. B) sGAG and collagen levels normalised to DNA show that cells within the self-organising microtissues produce significantly higher amounts of sGAG at day 21 and 28. C) sGAG and collagen levels normalised to wet-weight (ww) demonstrate near native levels of sGAG are accumulated in both groups by day 28, with significantly higher levels achieved when using microtissues. However, levels of collagen remain below native levels. N = 3, significant differences were determined using an ordinary two-way ANOVA with a Šídák’s multiple comparisons test where, * denote p < 0.05, ** denotes p < 0.01, *** denotes p < 0.001, and **** denotes p < 0.0001

While both self-organisation strategies supported a highly chondrogenic phenotype, histological analysis demonstrated clear differences between the two groups. A heterogeneity in cell and matrix phenotype, cellular morphology, and matrix deposition was observed through the depth of the single cell group. In addition, the cell single strategy generated a more spherically-shaped tissue, while the final shape of the tissue generated using the cartilage microspheres better mimicked the shape of the initial hydrogel mould. While the periphery of single cell constructs was rich in cartilage matrix components, its core stained weakly for sGAG and was diffusely mineralised. These features were evident as early as day 7 and persisted through the 28 day culture period. Cellular arrangement and morphology also changed through the depth of the single cell constructs, appearing hyaline-like in the construct periphery, but appearing more hypercellular and displaying an aberrant cobblestone morphology in the construct’s core (**Fig. 3A**). In contrast, the cartilage engineered *via* the bioassembly of cartilage microtissues appeared more homogeneous. After 14 days, the microtissues had undergone complete fusion, forming a unified macrotissue that stained positively for both sGAG and collagen. Unlike the single cell approach, homogeneous staining for sGAGs was observed by day 28. Moreover, cells within the microtissue group appeared to be round, and closely resembled native chondrocytes with some native-like stratification seen in the periphery of the tissue. Importantly, the ECM rich cartilage formed *via* microtissue self-organisation did not mineralise over 28 days of chondrogenic culture, indicative of a more stable hyaline cartilage phenotype (**Fig. 3B**). Collectively, these results provide compelling evidence for the formation of a superior and more homogeneous engineered cartilage through the bioassembly of microtissue precursors. Clear evidence of undesirable tissue heterogeneity and mineralisation was seen in cartilages engineered using single cells as biological building blocks, despite displaying other hallmarks of robust chondrogenesis.

**Figure 3.**
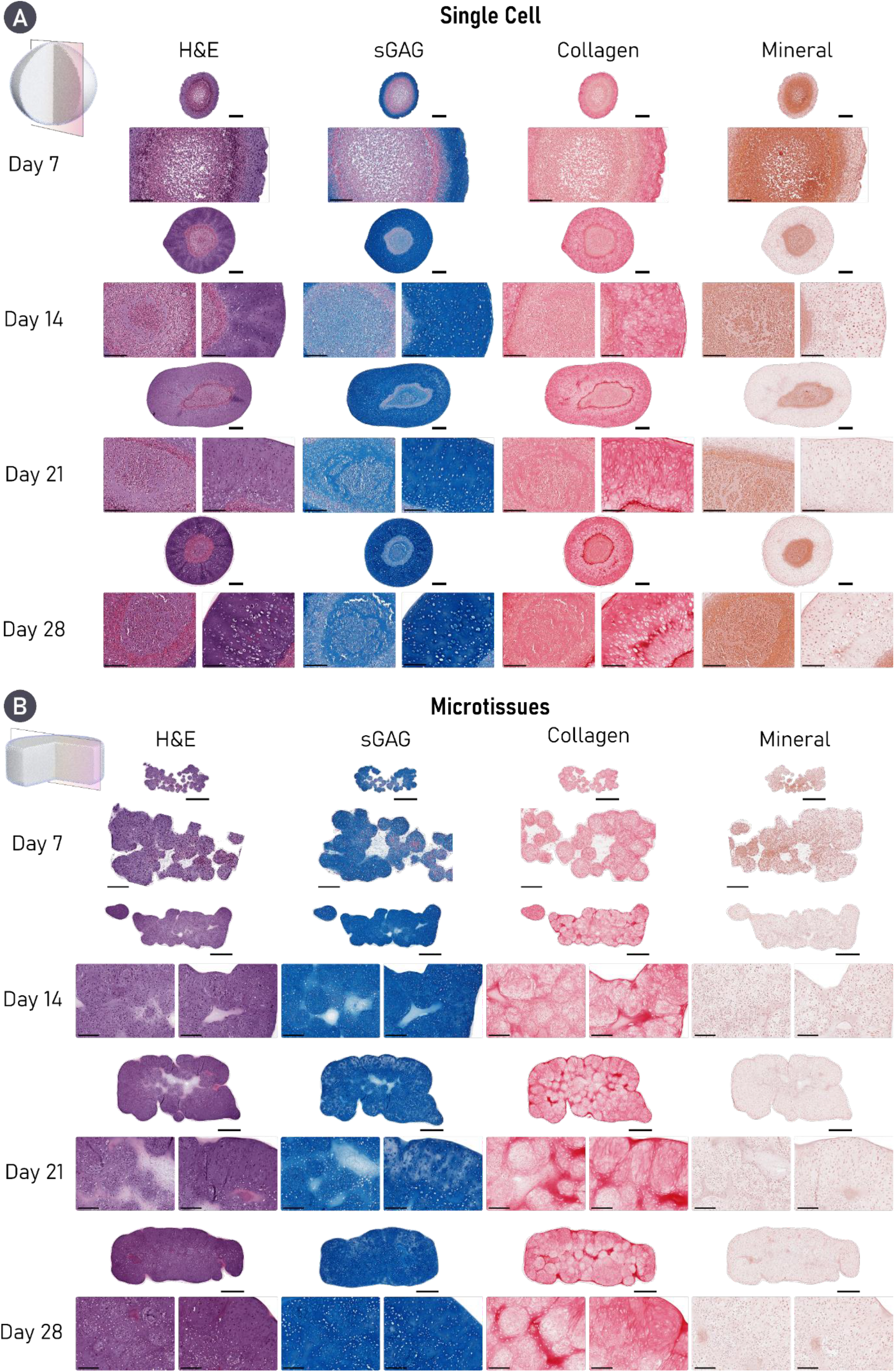
Cartilage microtissues self-organise a homogeneous matrix devoid of mineralisation. Histological analysis of engineered cartilage reveals robust accumulation of key cartilage ECM components in both groups, stained for using Alcian blue (sGAG) and Picrosirius Red Stain (Collagen). Alizarin red staining revealed evidence of cartilage mineralisation in tissues engineered using a single cell strategy. Additionally, a heterogeneous tissue structure and matrix deposition is clearly seen in the single cell group, whereas robust microtissue fusion results in homogeneous matrix deposition in the microtissue group by day 28. For both histological panels, an overview image is provided as well as zoomed sections for the core (left) and periphery (right) of the construct. A) Scale bars = 500 μm (Overview) and 200 μm (Zoom). B) Scale bars = 700 μm (Overview) and 200 μm (Zoom).

### Enzymatic treatment supports the development of a more biomimetic cartilage tissue

Since the use of similar enzymatic agents/regimes have shown great promise in enhancing the quality of engineered cartilages through modulation of the developing matrix [8,42,43], we next investigated the effect such an enzymatic regime would have on the self-organised tissues generated in this study. To this end, we exposed the engineered cartilages to a cABC solution for 4 hours at the mid-point (day 14) of chondrogenic cultivation. Unsurprisingly, biochemical evaluation of the engineered tissues at day 28 indicated that cABC treatment effectively reduced the total sGAG levels 2.48 fold and 2.96 fold in the single cell and microtissue groups respectively (**Fig. 4A**). There was still significantly higher sGAG/DNA in the cABC microtissue group compared to the cABC single cell group (**Fig. S1**). cABC treatment had no effect on total levels of DNA or collagen (**Fig. 4A, C & Fig. S1**). Despite having no effect on collagen synthesis, enzymatic treatment proved an effective method of altering the ratio of sGAG to collagen within the tissue towards a more collagen rich composition, typical of native AC (**Fig. 4B & C**). These findings support the use of cABC treatment as a strategy to significantly increase the relative amount of collagen within engineered cartilage (**Fig. 4C**).

**Figure 4.**
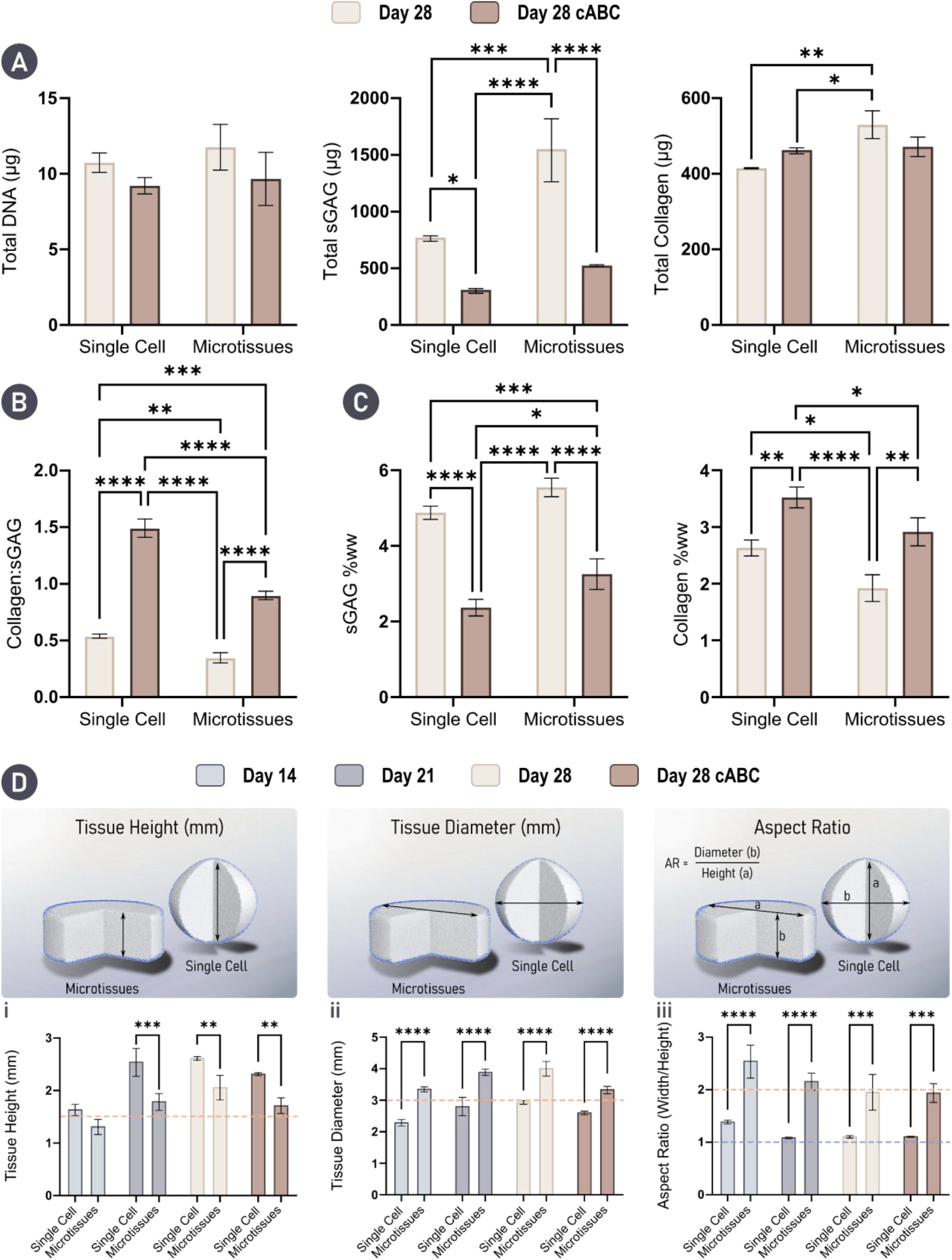
Enzymatic treatment results in a biomimetic ECM composition. Engineering with microtissues results in significantly better shape fidelity. A) Biochemical analysis of the engineered tissues indicated that cABC treatment did not affect total levels of DNA or collagen. However, enzymatic treatment did significantly decrease total levels of sGAG. B) Treatment using cABC significantly improved the collagen to sGAG ratio in both groups. C) sGAG and collagen levels as a percentage of tissue wet-weight (ww) show significant reduction in sGAG content with a concomitant significant increase in collagen content in both groups. Here, sGAG makes up a significantly larger portion of the total tissue wet-weight in the microtissue group compared to the single cell group following cABC treatment.). N = 3, significant differences were determined using an ordinary two-way ANOVA with a Tukey’s multiple comparisons test where, * denote p < 0.05, ** denotes p < 0.01, *** denotes p < 0.001, and **** denotes p < 0.0001 D) Physical characterisation including tissue: i) height, ii) diameter, and iii) aspect ratio indicates that superior control over tissue geometry is achievable by engineering cartilage macrotissues via the assembly of microtissues when compared to a single cell approach. Orange dashed lines denote the starting mould dimensions and aspect ratio. Blue dashed line denotes the aspect ratio of a sphere (AR = 1). N = 3, significant differences were determined using an ordinary two-way ANOVA with a Šídák’s multiple comparisons test where, * denote p < 0.05, ** denotes p < 0.01, *** denotes p < 0.001, and **** denotes p < 0.0001

In the context of engineering functional cartilage grafts, ensuring shape fidelity; that is to say the gross morphology of the final tissue closely matches the intended/designed geometry, as well as generating a mechanical competent graft are of great importance. To evaluate the physical properties of our engineered tissues, height, diameter and aspect ratio measurements were taken throughout the study (**Fig. 4D**). Collectively, these results indicated that by engineering cartilage grafts using microtissues as biological building-blocks yielded a final tissue with superior shape fidelity. Specifically, consistent vertical and lateral growth over 28 days of culture was observed, with engineered tissues maintaining the initial imparted cylindrical shape. In contrast, the use of single cells resulted in aggressive tissue contraction between days 14 and 21 (**Fig. 7Di**). Ultimately, the single cell approach yielded an engineered cartilage that was spherical, indicated by an aspect ratio of ~1 for days 21 and 28 (**Fig 4Diii**). Enzymatic treatment decreased the overall size (width & height) of the engineered tissues in both groups, but did not significantly impact the overall aspect ratio of the construct.

There was a trend towards increased mechanical properties with time in culture for the microtissue group (Note it was not possible to mechanically test the single cell constructs as they adopted a spherical shape in culture). Despite the dramatic loss of sGAGs with cABC treatment, there was also a trend towards increases in the Young’s and dynamic modulus with enzyme treatment. After 28 days in culture the compressive properties of the graft approached that of normal articular cartilage, with a Young’s modulus of 0.266 ± 0.157 MPa and a dynamic modulus of 0.821 ± 0.298 MPa for the enzymatically treated cartilages generated using microtissues (**Fig. S2**).

### Enzymatic treatment improves microtissue fusion and remodeling

We next sought to identify, histologically, how cABC treatment influenced matrix structure and composition. Predictably, sGAG staining in both cABC groups appeared weaker compared to the untreated controls, althought in line with the biochemical data, the microtissue group appeared to have recovered more sGAG with a less marked decrease in staining intensity (**Fig. 5A**). Despite not significantly increasing total collagen content (biochemically), cABC treatment resulted in a notably more intense collagen staining in both groups (**Fig. 5B**). As before, in the cABC single cell group a considerable portion of the engineered cartilage was composed of a core, devoid of sGAG and positively stained for mineral deposits (**Fig. 5C**). Consequently, the aforementioned densification of the collagen network was only observed in the periphery of tissue (**Fig. 5B**). It appeared that cABC treatment not only resulted in increased collagen density in the microtissue group, but also facilitated enhanced fusion and remodeling between the microtissue units. As a result of this, there was little to no evidence of their initially spherical geometry (**Fig. 5**). In both groups, cABC treatment appeared to increase mineralisation. However, this effect was seen to a greater extent in the single cell group, which exhibited diffuse mineral deposition throughout the tissue (**Fig. 5C**). In addition to creating a denser collagen network, cABC treatment also appeared to result in a smaller, rounder cellular morphology in the periphery of the single cell group and throught the tissue within the microtissue group (**Fig. 5 & Fig. S3**).

**Figure 5.**
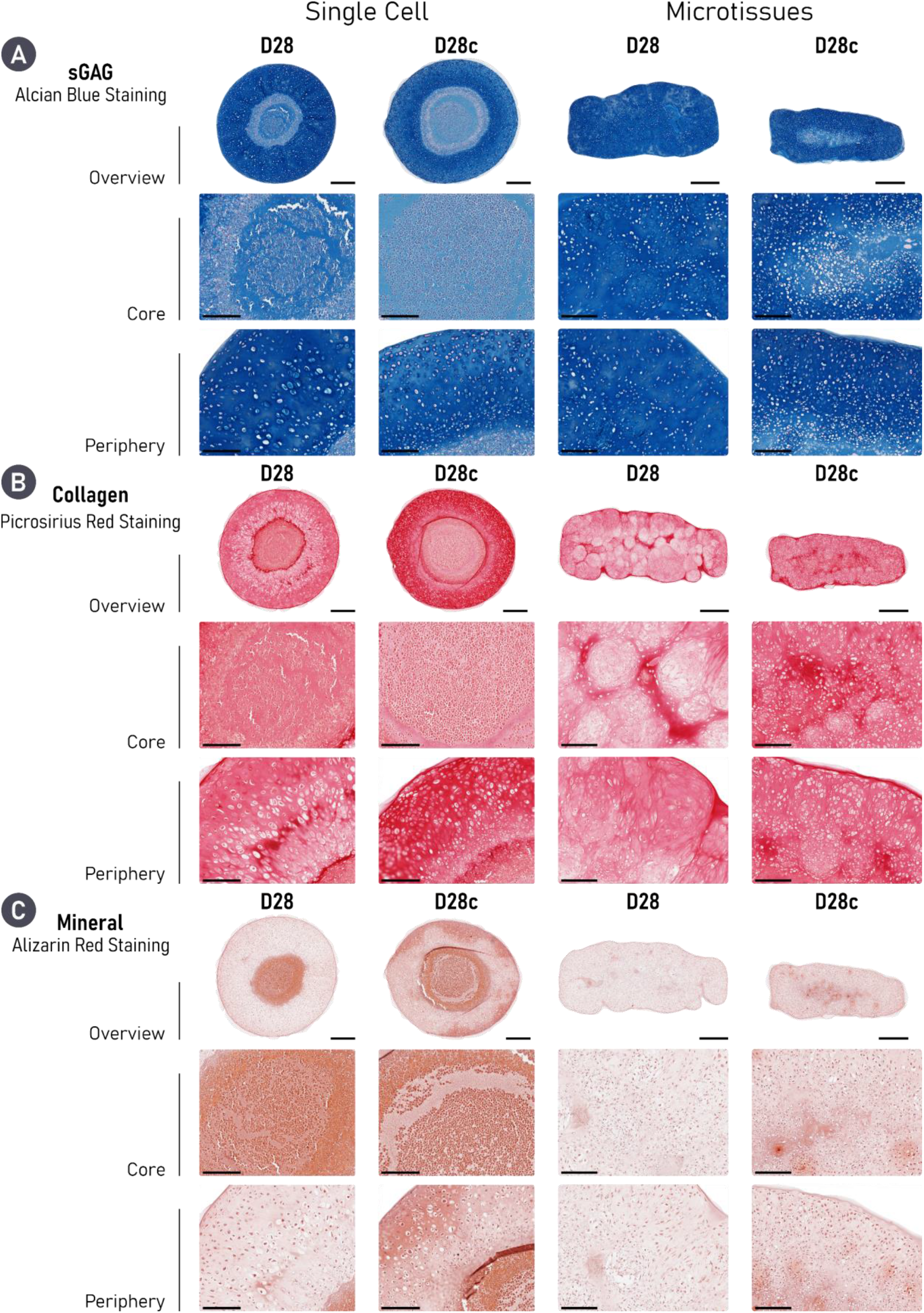
Enzymatic treatment results in densification of the collagen network and notably improves microtissue fusion. Histologically, both engineered cartilages treated with cABC exhibit robust chondrogenesis, as evidenced by diffuse positive staining for sGAG (Alcian Blue) and Collagen (Picrosirius Red Stain). cABC treatment causes a denser collagen network in both groups as evidenced by intense Picrosirius Red Staining. Importantly, enzymatic treatment appeared to significantly improve microtissue fusion. However, cABC also appeared to increase mineralisation when compared to untreated equivalents. Finally, treatment with cABC results in a smaller, rounder cell morphology in both groups. Scale bars: Single Cell = 500 μm (Overview) and 200 μm (Zoom). Microtissue = 700 μm (Overview) and 200 μm (Zoom).

### cABC treatment modulates the phenotype of self-organised cartilage

Having identified changes in ECM composition and structure following enzymatic treatment, we further investigated how removal of sGAGs during early tissue development influences tissue phenotype. To probe cartilage phenotype, immunohistochemical staining for collagen types I, II, and X were undertaken. All samples showed some positive staining for collagen type I, albeit weak, with the greatest expression in the untreated single cell group. In both groups, enzymatic treatment decreased the expression of collagen type I, with the most profound effect in the single cell group (**Fig. 6A**). All engineered cartilage stained intensely for collagen type II. Relatively homogeneous deposition for collagen type II was seen in both microtissue groups, whereas the core of the single cell groups expressed less collagen type II than the periphery, an outcome that was exaggerated by cABC treatment (**Fig. 6B**). Collagen type X was least expressed in the untreated single cell group, where it was predominantly found in the core of the tissue. In contrast, expression of collagen type X was noted in the periphery of the untreated microtissue group. In both groups, the use of cABC appeared to increase collagen type X deposition (**Fig. 6C**).

**Figure 6.**
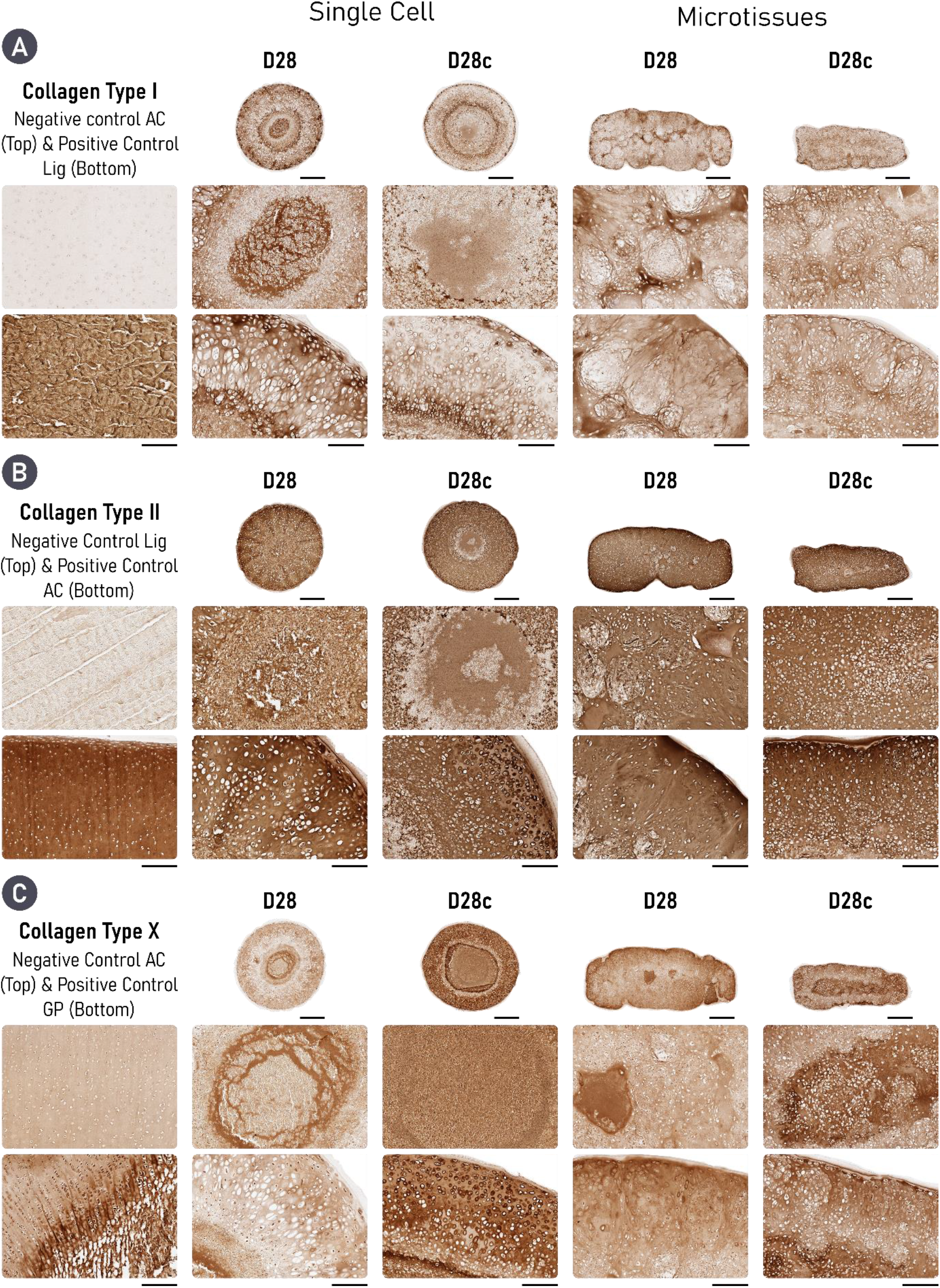
Cartilages engineered treated with cABC exhibit a less fibrocartilaginous, but more hypertrophic phenotype. A) In both self-organised strategies, cABC appears to reduce the expression of collagen type I. Equally, engineering cartilage via microtissue self-organisation also reduces the expression of collagen type I, with the most pronounced accumulation noted in the D28 single cell group. B) All groups diffusely express collagen types II and out of the three collagen sub-types it is the most abundant. C) Treatment of both engineered cartilages with cABC increase the expression of collagen type X. Scale bar = 700 μm (Overview) and 200 μm (Zoom)

### cABC alters collagen network organisation and supports the development of a zonally stratified tissue

Given the previous histological evidence of changes in the collagen network within the self-organised cartilage, polarised-light microscopy (PLM) was employed as a means of investigating how enzymatic treatment influences its spatial organisation. Untreated tissues assembled from single-cell building-blocks displayed localised organisation. However, as a result of their spherical gross morphology, the overall organisation of the collagen network did not closely match that of native AC (**Fig. S4**). Despite this, cABC treatment resulted in a clear change in the colour of the collagen fibres when viewed under polarised-light. This colour shift from green to orange/red is typically indicative of thickening of the collagen fibres, suggesting that cABC can be an effective method for increasing collagen fibre maturity (**Fig. S4**).

In the context of microtissue self-organisation, we identified that enzymatically treated tissues showed enhanced fusion between adjacent microtissues. When viewed under polarised-light, the engineered tissues appeared more homogeneous following cABC treatment. Additionally, a similar colour shift (from green to yellow/orange) in the collagen fibres could been seen following enzymatic treatment in the microtissue group (**Fig. 7A**). In the context of collagen network organisation, cartilages formed through microtissue self-organisation exhibited superior collagen stratification when compared to a single cell approach. Quantification of fibre orientation revealed that through enzymatic treatment, a more biomimetic collagen network was generated (**Fig. 7B, C & D**). Specifically, in both groups (untreated and cABC) the superficial and deep zones of the engineered tissue closely resembled native AC. However, the middle zone of the cABC treated microtissue group more closely matched the fibre orientation and distribution seen in native AC. Further quantification of the collagen fibre directionality within the engineered tissues demonstrated significant improvements through use of microtissues compared to single cells. The mean fibre orientation in the superficial zone was not statistically different to native AC for the microtissue groups. In the middle zone, only the cABC treated microtissue cartilage exhibited a mean fibre orientation that was not significantly different to native AC. In the deep zone of the tissue, all groups, including native AC, had a mean fibre orientation of ~90° (**Fig. 8A**). Fibre coherency was inferior in all engineered tissues compared to native AC. However, the highest levels of fibre coherency were seen in the tissues engineered using microtissues as biological building-blocks (**Fig 8B**). Collectively, PLM quantification revealed a superior tissue organisation *via* the self-organisation of microtissues compared to a single cell approach. Furthermore, enzymatic treatment during the self-organisation of the microtissues yielded a highly biomimetic collagen stratification that closely mimics native AC (**Fig. 8C & D**).

**Figure 7.**
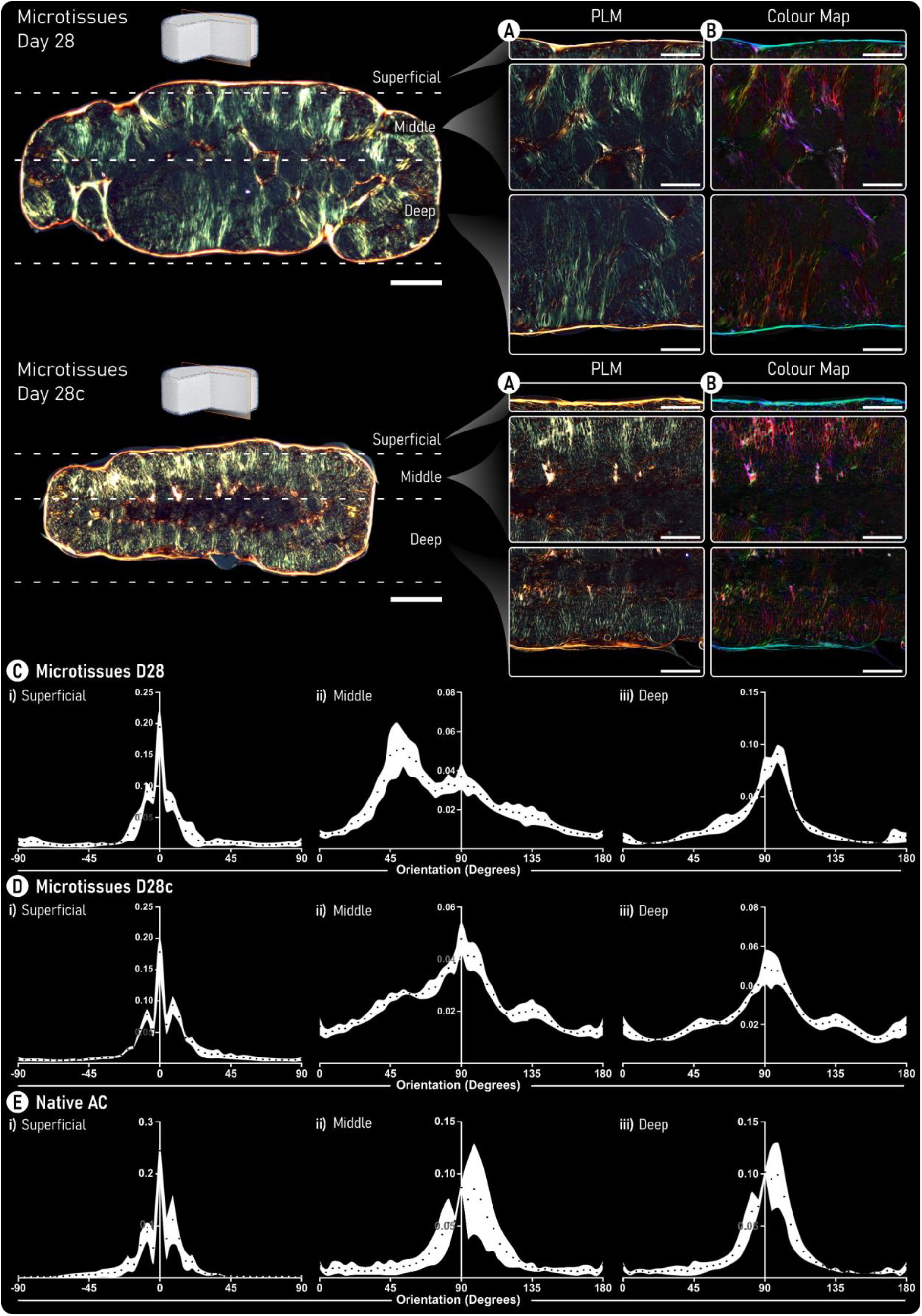
Enzymatic treatment increases collagen fibre thickness and supports the development of a biomimetic collagen network. A) Polarised-light microscopy (PLM) images of both untreated (top) and enzymatically treated (bottom) tissues generated by self-organization of cartilage microtissues. B) Colour maps generated from PLM images. Here, colour hue is used to indicate fibre orientation where, Red/Pink denotes fibres oriented at 90° and blue/cyan indicates fibres are oriented at 0°. (Scale Bars: Overview = 500 μm and Zoom = 250 μm). C & D) Quantification of the fibre orientation within the i) ‘superficial’, ii) ‘middle’, and iii) ‘deep’ zones of the engineered cartilage are provided for (C) untreated, (D) cABC treated engineered cartilage and (E) Native AC. Black data points represent the mean and the white area shows the standard deviation (n = 5).

**Figure 8.**
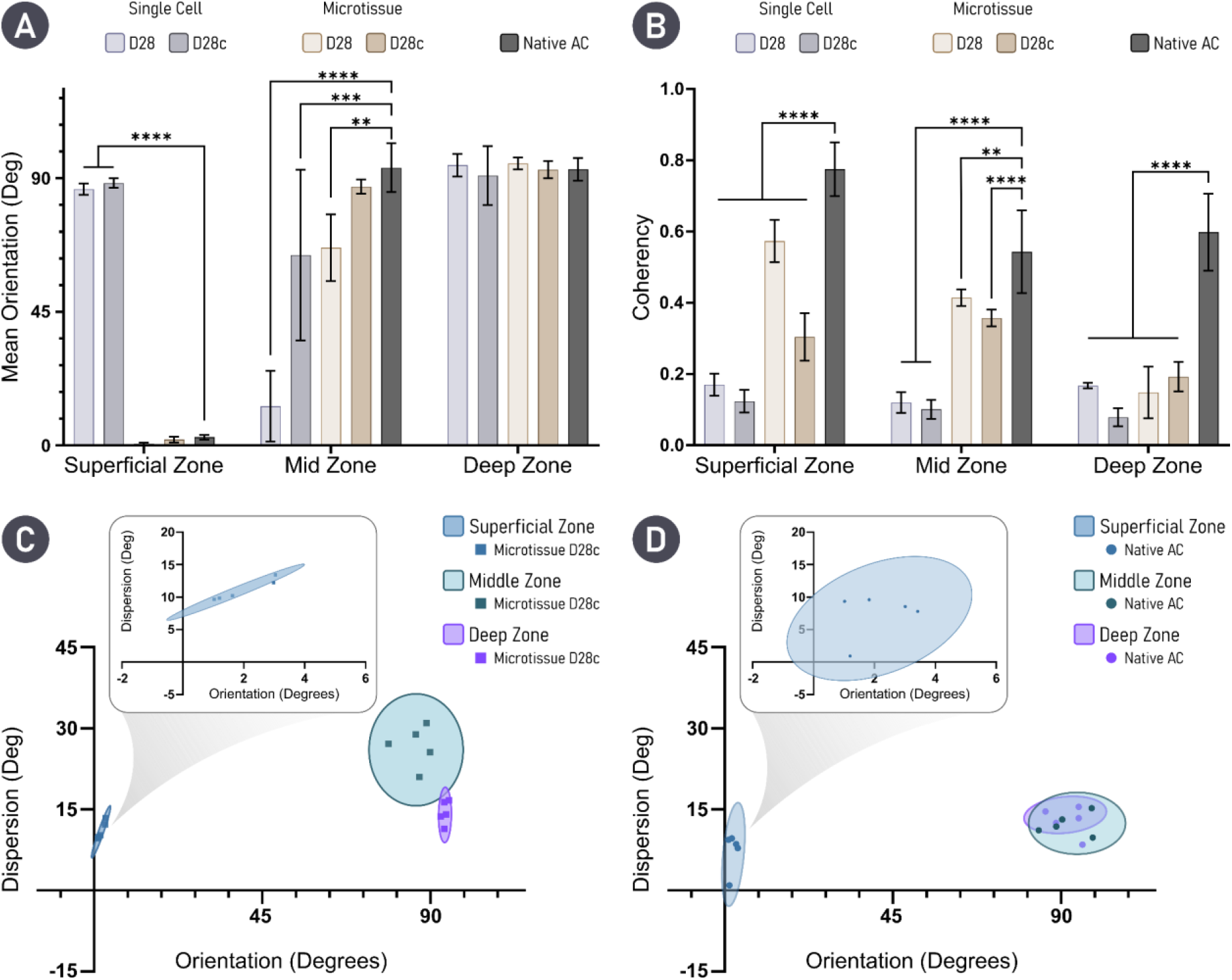
Quantification of collagen fibre orientation indicates that treatment with cABC supports the development of a biomimetic cartilage tissue. A) Mean average fibre orientation within the zones of the engineered tissues are compared with native AC. B) Fibre coherency; values approaching 1 indicate fibres are aligned in the same direction, while a value of 0 indicates dispersion of fibres in all directions. For engineered tissues n = 5 and native tissue n = 3. Statistical differences are determined using an Ordinary two-way ANOVA with a Dunnett’s multiple comparison test where the experimental groups are compared to the control of native AC. ** denotes p < 0.01, *** denotes p < 0.001, and **** denotes p < 0.0001. C & D) Mean orientation versus dispersion is given for cABC treated microtissue engineered cartilage and native AC respectively.

## Discussion

Here, we report a comprehensive comparison between the quality of engineered cartilage tissues formed *via* the self-organisation of single cells and microtissues. We demonstrated that both formats support robust chondrogenesis, however the cartilage engineered using microtissues as biological building-blocks exhibited a significantly richer ECM with higher biosynthetic output at a cellular level. This culminated in a neotissue with near native levels of sGAG accumulation after only 4 weeks of chondrogenic cultivation. Moreover, we demonstrate that the application of a catabolic enzyme (cABC) during tissue development aids in rebalancing the collagen:sGAG ratio within *in vitro* engineered cartilages, which are typically sGAG rich but collagen poor. Ultimately, cABC treatment promoted maturation of the collagen network within self-organised cartilage, significantly improved microtissue fusion, and resulted in a highly biomimetic collagen network with native-like zonal stratification.

In this study, we observed that employing a microtissue based strategy not only promoted superior cartilage ECM accumulation, but also supported the development of a more homogeneous graft. Specifically, a single cell approach supported noticeable heterogeneity within the engineered cartilage after 28 days of chondrogenic cultivation. Similar findings have been reported in the literature, whereby unfavourable radial heterogeneity in cell phenotype, matrix stratification, and occasionally the formation of a necrotic core has occurred following the chondrogenic induction of large, self-assembled cellular spheroids [46–50]. It seems likely that steep radial chemical and nutrient gradients readily develop within high density spheroidal cultures, leading to appropriate cellular differentiation and ECM accumulation at the circumference/periphery, but poor/off-target differentiation within the core of the implant. This phenomenon has also been observed in scaffold based approaches, where so called ‘core necrosis’ has impacted the *in vivo* therapeutic efficacy an engineered implants [51]. Latent forms of growth factors such as TGF-β have been employed in an attempt to alleviate some of the diffusion based limitations, by allowing the potent growth factor to penetrate the core of the engineered tissue prior to its activation and action [52,53]. Alternatively, as we have shown in this study, aggregate engineering is an effective method of generating tissues of scale through homogeneous cellular differentiation and ECM production. In agreement with our own findings, microtissue building-blocks have been used effectively to engineer millimetre scale tissues for cartilage [18,36,37,54], bone [13] and osteochondral [28,29,31] applications without obvious nutrient diffusion limitations.

In addition to overcoming diffusion limitations, engineering with microtissues also yielded a tissue with superior shape fidelity when compared to a single cell approach. It has been previously demonstrated that controlling tissue development and final geometry is achievable *via* self-organisation of MSCs through the use of bioprinting [38,55] as well as permanent [42,56–59] or transient cell adhesive substrates [60]. In this study, we found that MSCs seeded into a non-adhesive hydrogel well resulted in the formation of a large spherical aggregate, poorly suited as a tissue graft for biological joint resurfacing. This could be a result of increased cell-mediated contractions in MSC-only cultures. The use of protein-coated transwell membranes has previously been shown to allow the formation of cartilage discs rather than spheroids *via* the self-organisation of single cells [3,42,56–59]. However, the use of such modified membranes restricts the capacity to engineer user-defined/complex tissue geometries and can make the removal of the engineered tissue/graft difficult. As such, previous successful attempts of self-assembling single cells into a cylindrical construct in a non-adhesive hydrogel well have employed chondrocytes [11,42,43]. The data presented here and in literature suggests that engineering tissues with high shape-fidelity *via* the self-organisation of single cells is challenging and the success, at least in terms of shape fidelity, of an approach is closely coupled to the selected cell type. Our work indicates that using microtissues as biological building-blocks can significantly improve the control over final tissue geometry, and could provide a relatively simple platform for engineering patient specific grafts with user-defined geometries.

Although microtissue self-organisation enables the engineering of matrix-rich tissues at a millimetre scale with considerable control over macrotissue morphology, fusion between adjacent microtissues and their (re)modelling into grafts with biomimetic matrix organisation remains a challenge. Here we demonstrate that the temporal introduction of an exogenous remodelling enzyme, specifically chondroitinase-ABC (cABC), enhances microtissue fusion and macrotissue (re)modelling. Numerous different strategies have previously been employed to enhance the functional development of self-organised cartilage, including temporal [61] and spatiotemporal [56] exposure to physiological relevant growth factors [56,61], physical confinement [62], mechanical stimulation [63] and cABC treatment [8,42,43]. In line with our observations, previous studies have observed that the relative amount of collagen within engineered tissue significantly increases following cABC treatment [8,43]. This change in matrix composition correlated with significantly enhanced tensile mechanical properties without compromising compressive properties [8,43]. Through cleavage of sGAGs during early tissue development, we were also able to rebalance the collagen:sGAG ratio in the developing matrix towards more physiological levels. Interestingly, while temporal enzyme treatment lead to a dramatic reduction in sGAGs from the developing tissue, it did not negatively impact the compressive mechanical properties of the engineered graft. This can potentially be explained by the observed changes to the organisation of the engineered tissue, and specifically the development of a more biomimetic, arcade-like collagen structure which is known to play a key role in determining the mechanical properties of skeletally mature articular cartilage [64]. More significant improvements in tissue mechanical properties might be expected following multiple cABC treatments and/or extended culture periods to allow complete recovery in tissue sGAG content [42,43]. Other approaches that could be adopted in the future include the addition of an exogenous collagen crosslinking agent such as lysyl oxidase (LOX-2), in conjunction with cABC, to improve collagen fibril maturity and tissue functionality [8,41].

Temporal enzymatic treatment also resulted in significantly enhanced microtissue fusion as well as the formation of thicker, more organised collagen fibres. Others have reported similar findings whereby the addition of exogenous cABC during cartilage development has increased collagen fibril density and diameter [8,42]. Here, we leverage the colour shift in PLM as a qualitative method of determining an increases in fibril thickness/maturity [65,66]. Although this approach provided strong visual evidence that changes within the collagen fibril dimeter/maturity had occurred following enzymatic treatment, future studies will aim to leverage alternative techniques such as scanning electron microscopy as a means of quantifying the changes observed in fibril density and thickness. The removal of small regulatory matrix molecules, such as decorin, known to play a role in regulating collagen fibrillogenesis [67,68] has been proposed as a mechanism for how cABC treatment enhances collagen maturation [43]. Importantly, following cABC treatment, there was limited evidence of boundaries between adjacent microtissues, and it appears that employing this exogenous catabolic enzyme benefitted tissue (re)modelling. The observed structural and organisational changes to the collagen network could help to explain the encouraging trend towards increases in graft Young’s and dynamic modulus following temporal enzyme treatment. Although untreated cartilage microtissues were able to fuse together and form a continuous cartilage, evidence of their initial spherical geometry was still apparent after 28 days of culture. While it remains unclear what impact this ‘more disorganised’ ECM will have on implant functionality, PLM displayed clear improvements in the organisation of the collagen network within enzymatically treated cartilages which has implications in the broader field of aggregate based tissue engineering. The results of our study suggest that temporal enzyme treatment could be applied to a range of different tissue engineering strategies using cellular spheroids, microtissues or organoids as biological building blocks to fabricate scaled-up regenerative implants.

## Conclusion

This work has established a robust platform for engineering biomimetic cartilage tissues *via* cellular self-organisation. Fusion of microtissue building blocks generated a more hyaline-like cartilage tissue compared to a more traditional scaffold-free approach using individual cells. In addition, temporal exposure of the developing tissue to a remodelling enzyme (cABC) modulated matrix composition, enhanced microtissue fusion and tissue remodelling, and ultimately supported the formation of a denser, more mature collagen network that exhibited zonal organisation analogous to the native tissue. Our findings support the use of temporal enzymatic treatments when tissue engineering using multiple cellular spheroids, microtissues or organoids as a means of generating more functional grafts.

## Acknowledgements

This publication was developed with the financial support of Science Foundation Ireland (SFI) under grant number 12/RC/2278 and 17/SP/4721. This research is co-funded by the European Regional Development Fund and SFI under Ireland’s European Structural and Investment Fund. This research has been co-funded by Johnson & Johnson 3D Printing Innovation & Customer Solutions, Johnson & Johnson Services Inc.

