## Supporting information for "Temporal enzymatic treatment to enhance the remodelling of multiple cartilage microtissues into a structurally organised tissue"

* Corresponding Author

### Supplementary Information

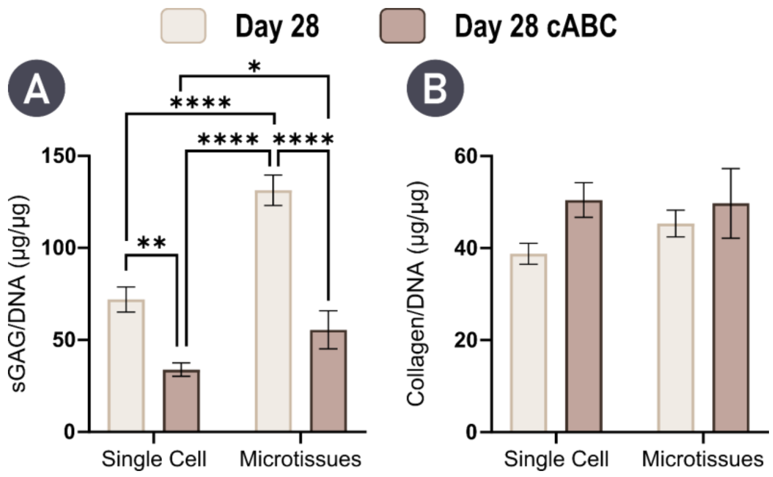


***Figure S1. Microtissue self-organisation results in higher biosynthetic output for sGAG following cABC treatment.*** *A) sGAG content normalised to DNA levels in engineered cartilages after 28 days indicated significantly higher biosynthesis of sGAG in the microtissue group following enzymatic treatment. B) Collagen content normalised to DNA shows trends towards increased production following enzymatic treatment, but the differences are not statistically significant. N = 3, significant differences were determined using an ordinary two-way ANOVA with a Tukey's multiple comparisons test where, * denote p < 0.05, ** denotes p < 0.01, *** denotes p < 0.001, and **** denotes p < 0.0001*

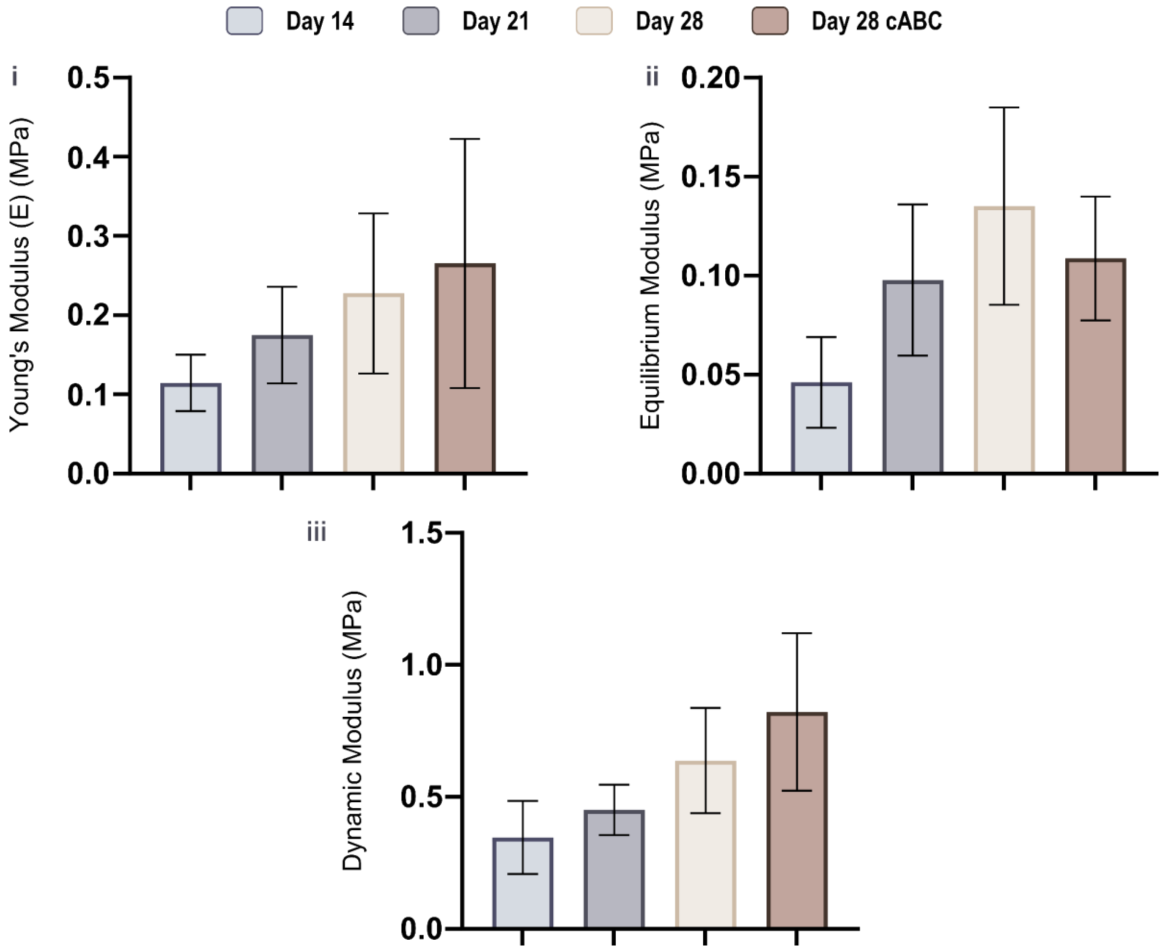


***Figure S2. Mechanical properties of self-organised cartilage formed via microtissues shows trends towards increases following enzymatic treatment. A****) Mechanical characterisation of cartilage tissues engineered using microtissue building blocks show the biofabrication of a mechanically competent tissues and trends towards improved mechanical properties with enzymatic treatment (N = 3).*

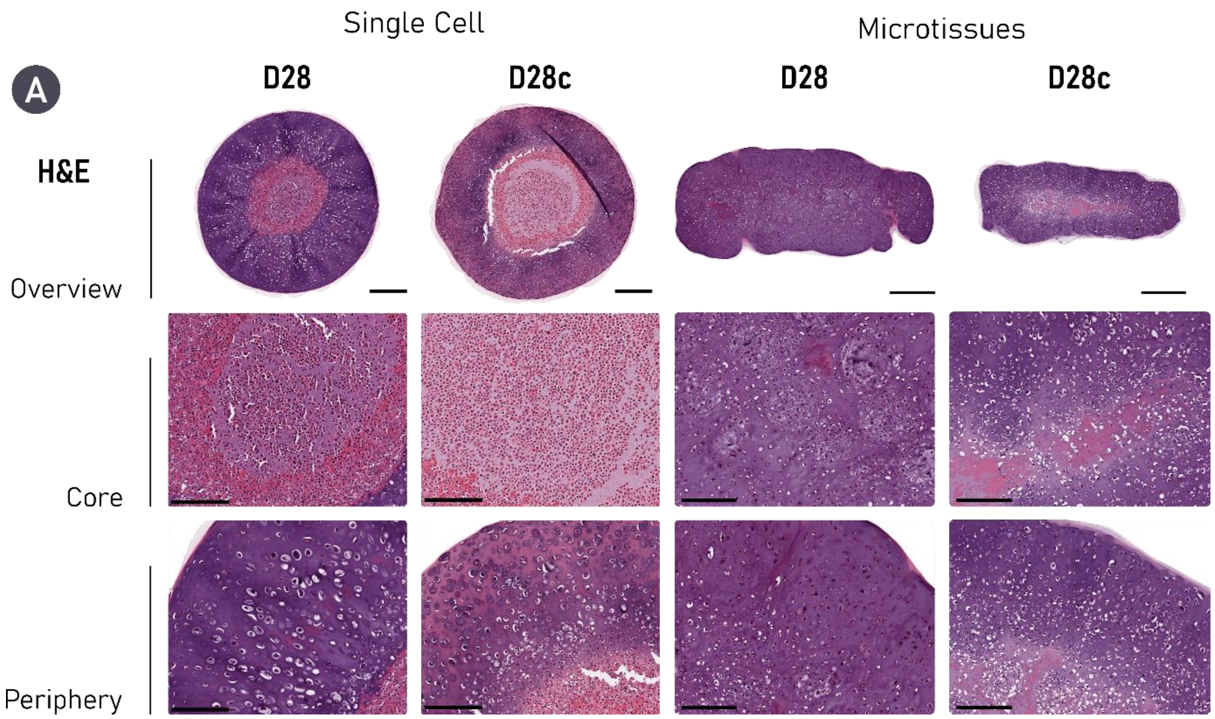


***Figure S3. Haematoxylin & Eosin Staining for untreated and enzymatically treated self-organised cartilages at Day 28.*** *Scale bars: Single Cell = 500 µm (Overview) and 200 µm (Zoom). Microtissue = 700 µm (Overview) and 200 µm (Zoom).*

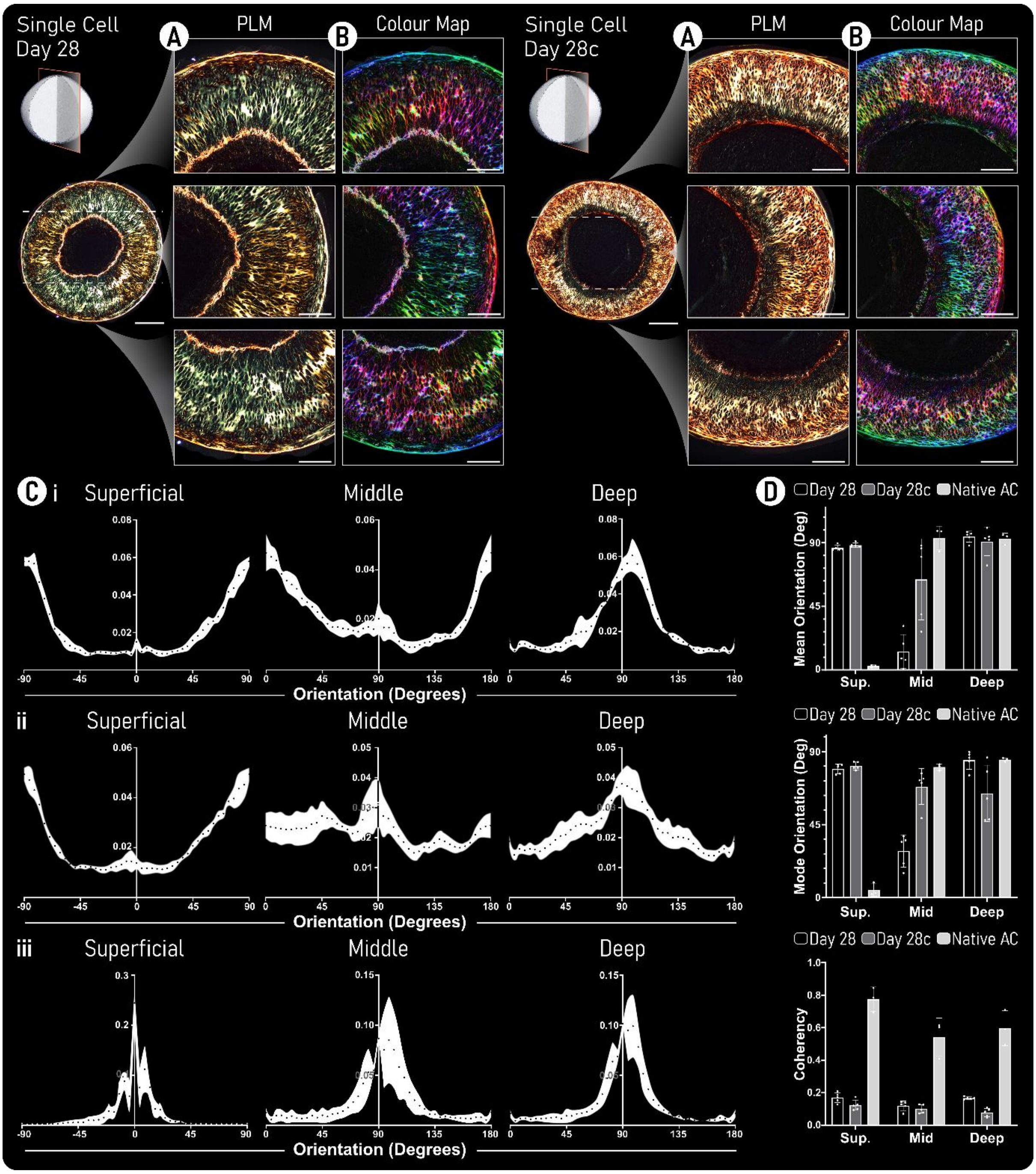


***Supporting Figure 4. Polarised-light microscopy reveals a single cell self-organisation approach does not yield a biomimetic collagen organisation. However, cABC treatment results in thicker collagen fibres.*** *A) For both untreated (left) and enzymatically treated (right) polarised-light microscopy (PLM) images are provided. B) Colour maps generated from these PLM images are given. Here, colour hue is used to indicate fibre orientation where, Red/Pink denotes fibres oriented at 90° and blue/cyan indicates fibres are oriented at 0°. (Scale Bars: Overview = 500 µm and Zoom = 250 µm). C) Quantification of the fibre orientation within the 'superficial', 'middle', and 'deep' zones of the engineered cartilage are provided for untreated (i), cABC treated (ii), and native AC (iii). Black data points represent the mean and the white area shows the standard deviation. D) Within the same three tissue zones, the mean average orientation, mode average orientation, and coherency are provided. For the latter, values approaching 1 indicate fibres are aligned in the same direction, while a value of 0 indicates dispersion of fibres in all directions. For engineered tissues n = 5 and native tissue n = 3.*
